# Enhanced neuroimaging with a calcium sensor in the live adult *Drosophila Melanogaster* brain using closed-loop adaptive optics light-sheet microscopy

**DOI:** 10.1101/2023.01.06.522997

**Authors:** Antoine Hubert, Georges Farkouh, Fabrice Harms, Cynthia Veilly, Sophia Imperato, Mathias Mercier, Vincent Loriette, François Rouyer, Alexandra Fragola

## Abstract

We report on an Adaptive Optics (AO) Light-Sheet Fluorescence Microscope compatible with neuroimaging, based on direct wavefront sensing without the requirement of a guide star. We demonstrate fast AO correction, typically within 500ms, of in-depth aberrations of the live adult *Drosophila* brain, enabling to double the contrast when imaging with structural or calcium sensors. We quantify the gain in terms of image quality on multiply neuronal structures part of the sleep network in the *Drosophila* brain, at various depths, and discuss the optimization of key parameters driving AO such as the number of corrected modes and the photon budget. We present a first design of a compact AO add-on that is compatible with integration into most of reported Light-Sheet setups and neuroimaging.

## 1 Introduction

Understanding the functional organization and connection of the brain is currently a huge challenge in the field of neuroscience, which is of crucial importance to understand how the brain functions and the mechanisms behind the different brain disorders, for example, the neurodegenerative diseases like Alzheimer’s and Parkinson’s diseases. In particular, such a challenge requires to decode the functional behavior of neuronal networks over larges scales. To this end, functional neuroimaging is a key step and recent developments around calcium and voltage sensors now enable the spatio-temporal monitoring of functional connections between neurons in living samples. In order to map the brain activity in specific areas while remaining as minimally invasive as possible, many strategies have been implemented over the last years, targeting the right combination of sensitivity, resolution and speed [1, 2, 3, 4]. Among these strategies, Light-Sheet Fluorescence Microscopy (LSFM) [5] is a powerful imaging technique that allows both structural and functional imaging of neuronal networks, with appropriate spatio-temporal capabilities and minimal phototoxicity, making it particularly suitable for imaging living organisms [6].

The intrinsic optical sectioning of LSFM enables to image with high Signal-to-Background Ratio (SBR), over a large Field-of-View (FOV) using modern objectives, which also preserves a high spatial resolution at the single cell level. Starting from initial setups, various types of LSFM approaches have been reported [7, 8, 9], including improvements at the excitation path to provide a thin beam over a large FOV [10, 11, 12], and advanced setups mostly based on remote focusing enabling fast 3D imaging [13, 14, 15]. These new setups make LSFM undeniably one of the most recommended optical sectioning methods for high-speed imaging of 3D samples. Functional imaging using LSFM has therefore been widely used in various samples, such as zebrafish, *C. elegans* and *Drosophila melanogaster*, but suffers from the drastic reduction of SBR encountered in depth due to the appearance of strong optical aberrations [16]. In order to compensate for these aberrations, Adaptive Optics (AO) has been implemented on several LSFM setups, in both the excitation and the emission paths. Such implementations mainly differ on how aberrations are measured, either using sensorless methods [17], or using direct wavefront sensing with a guide-star such as a fluorescent bead [18] or a virtual point source taking advantage of the confinement of multiphoton excited fluorescence and using a scan/descan geometry [19]. These methods demonstrated significant improvement in SBR and in-depth resolution, but at the cost of either low temporal performance inducing phototoxicity for sensorless approaches, invasiveness when using beads, or implementation complexity and cost when based on multiphoton excitation.

Recently, we demonstrated AO-LSFM based on direct wavefront sensing, taking advantage of the use of an Extended-Scene Shack-Hartmann (ESSH) wavefront sensor [20]. The key benefits of the approach are its temporal performance and accuracy when compared to sensorless methods, the non-necessity of a guide-star, but also its minimized instrumental complexity when compared to non-linear excitation approaches. This strategy promises easy AO implementation on LSFM setups, fast closed-loop correction for minimal phototoxicity and easy use, and is compatible with a design as an add-on to existing systems. As a first proof of concept, after having characterized the metrological performances of the sensor, we reported significant improvement of image quality in depth in the live *Drosophila* brain, targeting neurons of the sleep network [20]. However, these initial results were obtained for structural imaging, and no demonstration of the capability of the approach to enhance functional imaging was performed.

In this paper, we demonstrate the capability of this new AO-LSFM approach to perform enhanced neuroimaging in freshly dissected samples, at significant depths, with significant gain for structural or calcium signals thanks to aberrations correction. In particular, we present an enhanced setup providing minimal toxicity induced by the AO process, no consumption of photons dedicated to neuroimaging, and improved speed of the AO loop. This enhancement is performed through a combination of an instrumental upgrade and of a minimal requirement in terms of sample preparation. We report on structural and calcium images in depth of the live, adult *Drosophila* brain with a gain of at least 2 in terms of signal and a significant gain in terms of resolution, both brought by AO. Finally, we provide an analysis of a key parameter driving the AO process and performance, such as the number of modes, and provide initial analysis and discussion of the isoplanetic zone in the adult Drosophila brain in the context of our adaptive optics approach.

## 2 Methods

### 2.1 Two-color labeling and spectral separation for photon budget optimization

When targeting functional imaging, it is of critical importance that the photons corresponding to the functional signal of interest are directed to the imaging camera with minimal loss, since this signal is usually close to the sensitivity of detectors. Moreover, the variation of the functional signal would be an additional difficulty if it were used to measure aberrations. In our initial proof of concept for structural imaging [20], fluorescence photons were shared equally, using a 50:50 beamsplitter, between the imaging path and the wavefront sensing path for the sake of experimental simplicity. Jumping to functional imaging requires to take action on the optimization of the photon budget for both optical paths. In this new setup, we demonstrate the combined use of a specific sample preparation and of a dual - or more - color optical setup at the emission of the LSFM to use photons which are continuously emitted in a specific spectral range for the AO process, while photons corresponding to the calcium signal within a second spectral range are directed to the imaging camera with negligible loss. The principle is the following:

- The genetically-encoded reporters are modified such that a specific labeling is available for structural imaging with a given fluorescence emission spectrum, while another labeling is used for functional imaging with a second fluorescence spectrum, the 2 emission spectra being separated by a shift compatible with efficient spectral separation.
- The optical setup is modified such that it provides the excitation wavelengths corresponding to the 2 fluorescent labels, and the capability to separate the emission spectra with minimal photon loss.

While the first step could be seen as a constraint, current genetic methods now perform such sample preparation in routine, for multiple animal models used in neuroimaging. The addition of a stable structural fluorescence signal to the functional imaging has also the benefit of properly locating the functional signals [4] and has been used as reference to compare the activity level of neurons between different animal samples [21]. Moreover, structural labelling usually provides continuous fluorescence signal, thus enabling constant feed of photons to the wavefront sensor that is driving the AO process. Details about the adapted setup and corresponding sample preparation are given in the next paragraphs.

### 2.2 Optical setup

The optical setup is based on the previously reported AO-LSFM system, for which full implementation details are described in [20], in particular regarding the ESSH wavefront sensor design and metrology performance. Fig.1(left) provides a simplified representation of the setup (conjugation lenses are not represented), focusing on modifications required for efficient functional imaging (see previous section). In the excitation path, two lasers (Cobolt 06-MLD-488nm and Cobolt Jive-561nm) are coupled inside a monomode fiber, providing adequate excitation wavelengths for the green fluorescent protein (GFP) and its derived variant for calcium imaging GCaMP7b [22], and the red fluorescent protein ChRFP, (see next section – sample preparation). A 5μm thick light-sheet is formed using a galvanometric mirror (GVS002, Thorlabs) and sent to the sample through an air objective (10x, NA=0.3, Olympus) and a custom sample holder with No.1.5 coverslips on its sides. A variable diameter aperture diaphragm placed in the back focal plane of the illumination objective allows to adjust the thickness of the light-sheet. Since the groups of neurons of interest in the present study, involved in the circadian cycle of the adult *Drosophila*, do not cover the whole imaging field of view of the imaging camera (see section 4. Results), we adjusted the aperture diaphragm in order to decrease the light-sheet thickness down to 5μm when compared to our previous publication (7μm). This allowed to improve optical sectioning and signal-to-background ratio of achieved images. Emitted fluorescence is collected through a high-NA water immersion objective (25x, NA=0.95, Leica). Fluorescence emitted by GFP (for test experiments) or GCaMP (for functional imaging) is shown in green in Fig.1(left) and is sent to an sCMOS camera (ORCA Flash V2 - Hamamatsu), while fluorescence corresponding to structural ChRFP labeling (in red in Fig.1(left)) is sent to the ESSH wavefront sensor (custom HASO design from [20] – Imagine Optic). Fluorescence signals are spectrally separated using a combination of a dichroic filter (Semrock FF556-SDi01) and of bandpass filters (Chroma ET525/50, Semrock FF01-618/50). Before reaching the spectral separation unit, fluorescence signal is reflected by a high stroke, high linearity Deformable Mirror (DM, Mirao 52e – Imagine Optic) for aberration correction. The Field of View (FOV) is 350×350μm^2^ on the camera and each microlens form a 130×130μm^2^ FOV image on the ESSH sensor. Before performing imaging experiments, the static system aberrations were calibrated by measuring the aberrations from a point source, coming from a sample made of sparse 2μm beads in a 2% agarose gel at the sample location, using the same direct wavefront sensor with a centroïd algorithm. Then a closed-loop was applied to determine the deformable mirror shape that compensates for these aberrations. Finally, optimization was achieved on the image of a bead in order to take into account the non-common path aberrations coming from the spectral separation unit and visualize a diffraction-limited Airy disc coming from the bead (see Fig.S1 for additional information). For this final shape of the deformable mirror, the corresponding wavefront is measured and recorded to be used as a reference for the ESSH in further AO experiments. The deformable mirror shape is recorded as the instrument-corrected shape, to be applied before any imaging. All results presented in this paper without AO are instrument-corrected so that the reported gain brought by AO only comes from the correction of sample induced aberrations. The wavefront calculation procedure has been described in detail in a previous article [23], and the slopes calculation is based on a Southwell geometry. The slopes calculation process based on intercorrelations follow the pseudo-code described in Anugu et al [24].

**Figure 1:**
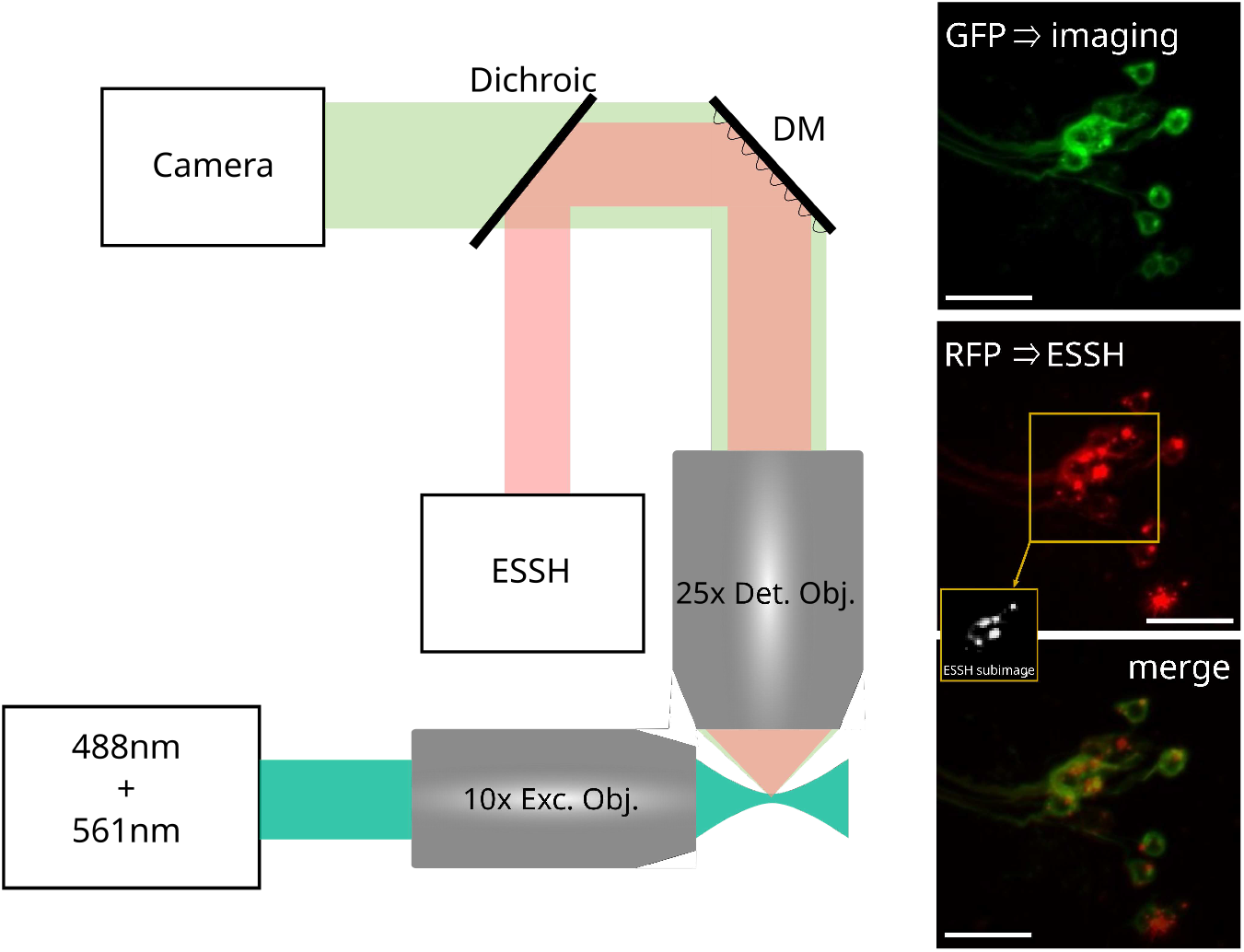
(left) AO-LSFM schematic setup for a functional imaging, as modified from [20]. Red and green beams are of different sizes for representation purpose only (right) Example of cell bodies of FSB neurons in the *Drosophila* brain with GFP and ChRFP double labeling. Images acquired on a conventional epifluorescence microscope (Zeiss AxioImager M2). (top) GFP signal used for imaging. (center) ChRFP signal used to perform wavefront sensing and real sub-image associate on the ESSH. (bottom) Both previous signals merged, the localization is identical. Scale bar: 100μm

### 2.3 Sample preparation

Samples consist of freshly dissected adult *Drosophila* brains with sleep-promoting neurons of the fan-shaped body (FSB) expressing either GFP or GCaMP7b, and ChRFP fluorescent proteins [23E10-GAL4 > UAS-mCD8::GFP, UAS-mCD8.ChRFP or 23E10-GAL4>UAS-GCaMP7b, UAS-mCD8.ChRFP], without any sample processing such as sample fixation or clarification. The emission spectra of the green and red fluorescent proteins are respectively centered at 509nm and 610nm. It corresponds to a spectral separation of 101nm, which ensures negligible impact of the wavelength dependency of aberrations of our samples [25] on the quality of the AO correction. Moreover, the width of the emission spectra of the fluorescent proteins, combined with the position of the emission peak, allows a minimal loss of functional GCaMP7b signal when passing through the chosen dichroic filter.

The following genetic lines of *Drosophila* flies were obtained from Bloomington Drosophila Stock Center: 23E10-GAL4 (#49032), UAS-mCD8::GFP (#32186), UAS-mCD8.ChRFP (#27391), UAS-GCaMP7b (#80907). Standard genetic crosses were used to obtain flies expressing the dual green and red fluorescent proteins in the same neuronal structures, so that the correction of aberrations based on the structural signal is optimized for the functional signal. More precisely, if red and green labeling were not done on the same structures, wavefront measurement based on the red structural signal could lead to a different result than a wavefront measurement based on the green functional signal in the case of large aberrations strongly limiting the isoplanetic patch down to a smaller size than the field of view captured by one microlens. Fig.1(right) illustrates the achieved sample preparation: it shows the FSB neurons as imaged at 30-55 μm depth using a widefield microscope for GFP (left), ChRFP (middle) and numerically merged signals (right), over a 350 × 350 μm^2^ FOV. The thumbnail on the middle image corresponds to the area of the image captured by each microlens of the ESSH in order to perform wavefront sensing on an extended scene (see [20] for details of the wavefront sensing process). FSB neurons project axons into a denser area at about 60-80μm depth; we also report AO-enhanced images of such neurons later in the manuscript (see 3.1).

### 2.4 Number of corrected modes

The choice of the number of aberrations modes to be corrected in the AO process is critical to achieve the best quality of aberration correction while still ensuring the capability of the AO loop to properly converge. Spherical aberration is one major aberration to be compensated for in biological samples, in particular due to the index mismatch with its surrounding medium in microscopy observation, but its sole correction does not ensure optimal image quality [26, 27]. Correcting higher order modes provides better performance. However, taking into account high order modes, corresponding to high order DM eigenmodes of the interaction matrix from the AO calibration process, induces higher sensitivity to the WF measurement noise, thus higher probability of a divergence of the AO loop [28]. Even though such a trade-off regarding the number of DM eigenmodes to be taken into account in the AO process has been widely reported as crucial, there has been poor experimental demonstration of its impact on AO-enhanced microscopy in biological samples. We thus experimentally determined such a trade-off in our setup. With the chosen DM, 52 eigenmodes are accessible for the correction, already corresponding to very high WF spatial frequencies such as defined e.g. by Zernike polynomials. We compared on two *Drosophila* samples, prepared according to the protocol described in the previous section, images of neuronal projections exhibiting high spatial frequencies without AO and with AO with different numbers of DM eigenmodes. When keeping all 52 modes, the AO loop starts diverging after approximately 10 iterations for a high gain value of the AO feedback loop (0.8). Fig.2 presents a comparison between no AO, AO using 16 and 36 DM eigenmodes, on the same FOV, 60 to 80μm deep, for 2 different areas of neuronal projections. For each case, a representation of measured aberrations is given by the values of the first 15 Zernike coefficients, up to the 5th order spherical aberration. The Zernike coefficients beyond mode 15 are negligible and are therefore not shown here. Without AO, measured aberrations (Fig. 2, left) confirm results from previous publications [19, 26], i.e. 3rd order spherical aberration (8th Zernike coefficient) being dominant but accompanied by mostly 3rd order aberrations. AO correction based on 16 DM eigenmodes (Fig. 2, middle) show strongly reduced Zernike values, but only provides a moderate gain regarding image quality. Interestingly, 5th order spherical aberration (15th Zernike coefficient) shows in this case an increased value after AO (confirmed by the corresponding wavefront map shown in Fig. S2), that is likely to explain the limited visual gain on images. We attribute this behavior to the fact that the DM eigenmodes do not correspond to Zernike polynomials. DM eigenmodes are indeed linked to the membrane shape and the spatial distribution of actuators, and even in the case of the Mirao 52e DM exhibiting a circular membrane with regularly distributed actuators underneath, there is no single DM eigenmode perfectly describing the 5th order spherical aberration, this discrepancy increasing with the order of the aberration to be corrected. For the DM to correctly describe the 5th order spherical aberration, a combination of several DM eigenmodes, including high orders, are required. As spherical aberration of various orders is very present in depth, the restriction of the number of DM eigenmodes to 16 strongly limits the capability of the AO loop to compensate the 5th order spherical aberration. This hypothesis is confirmed by the third experiment (Fig. 2, right), showing the result of AO correction using 36 DM eigenmodes. In this case, there is a visual increase of image quality, regarding image contrast, and all first 15 Zernike coefficients including the 5th order spherical aberration are reduced to values close to zero. This improvement is confirmed quantitatively by line profiles presented in Fig. S2 (supplementary material). For the two 2 experiments involving AO, convergence of the loop was ensured after 5-6 iterations, and stability was properly maintained afterwards.

**Figure 2:**
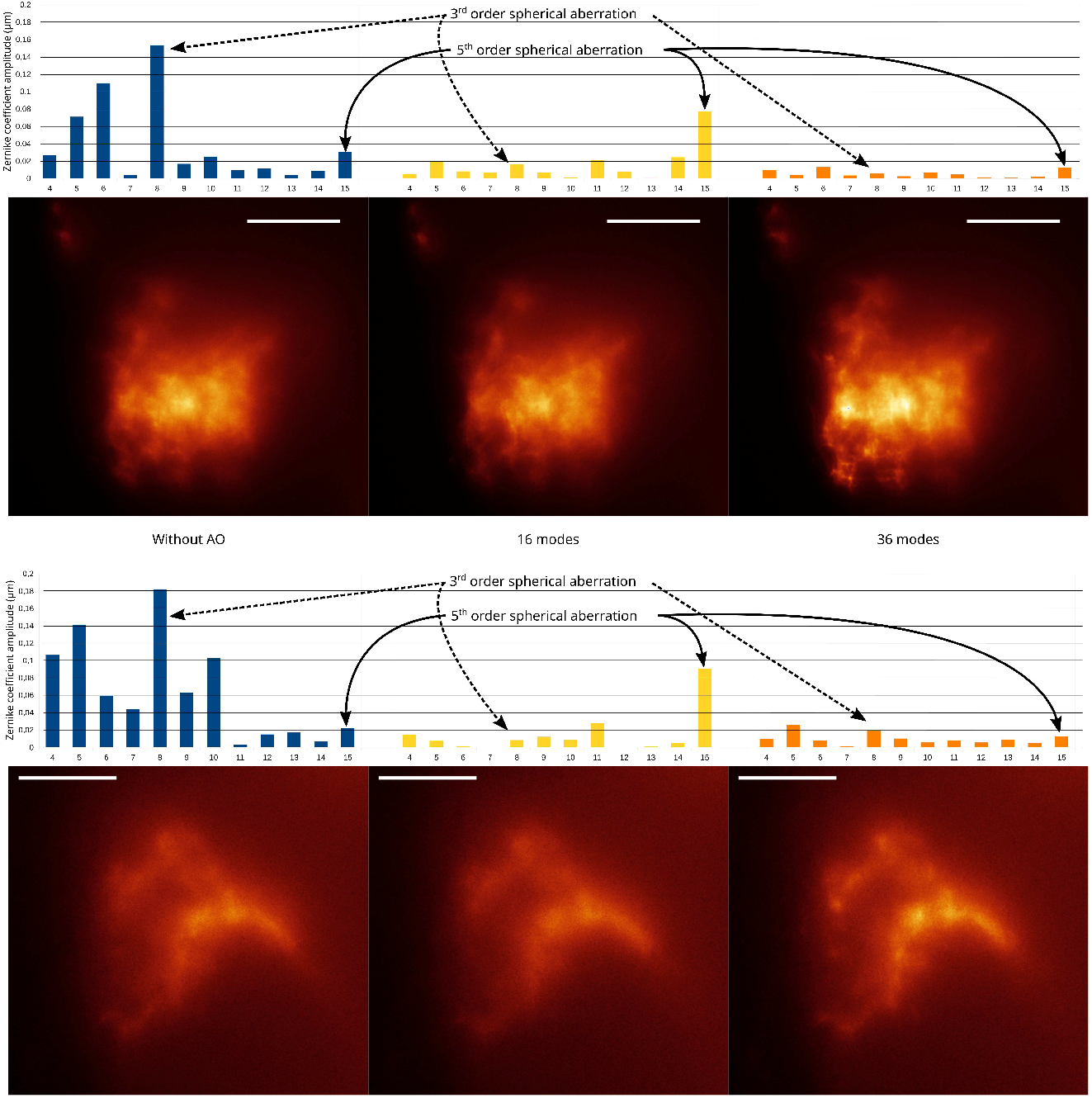
Images of neuronal projections of FSB neurons between 60-80μm deep in two different freshly dissected *Drosophila* brains without AO (left) and with AO with 16 modes (middle) or 36 modes (right). Scale bar 20μm. Graphs represent absolut value of Zernike modes from 4 (Astigmatism 0) to 15 (5th order spherical aberration) corresponding to images. Associated wavefront and full FOV for top image is presented in Fig.S2

## 3 Results

### 3.1 Ex vivo quantification of AO performance

We first demonstrate the performance of our new AO-LSFM setup on *ex-vivo*, freshly dissected adult *Drosophila* brain samples, with a structural dual color labeling and prepared according to the previously described procedure. Neuron cell bodies (Fig. 3(a)) and neuronal projections (Fig. 3(b)) were imaged at respective approximate depths of 30-55 and 60-80μm. Exposure times were 200ms on the imaging camera (GFP) and 100ms on the ESSH wavefront sensor (ChRFP). For each structure of interest in (a) and (b), Fig.3 shows images before (left) and after (right) AO correction, demonstrating a strong gain in image quality. More specifically, sub-images show the corresponding measured wavefronts, that drop from 256nm to 34nm root mean square (RMS) in Fig. 3(a) and from 377nm to 37nm RMS in Fig. 3(b), which in both cases correspond to a correction better than λ/15 RMS as compared to the Maréchal criterion defining diffraction-limited imaging at λ/14 RMS residual wavefront error. From the measured wavefronts, it can be observed that, as expected, spherical aberration is dominant – together with astigmatism – before the AO process. In order to quantify the gain brought by AO, we computed the average radial Fourier transform for each image (top graph in the middle of Fig. 3(a) and Fig. 3(b)): it shows a clear enhancement of medium spatial frequencies up to 50% in Fig. 3(a) and 30% in Fig. 3(b). This enhancement is quantitatively confirmed by profiles corresponding to white arrows in Fig. 3(a) and Fig. 3(b) (bottom graph in the middle). Profiles show a clear contrast enhancement by a factor of more than 2 in both cases, and a corresponding increase in terms of spatial resolution. The latter is particularly visible in Fig. 3(a): since GFP corresponds to a membrane labelling, AO provides a better view of the nucleus that corresponds to a dip in the profile (and a dark, round structure in the middle of the cytoplasm in the fluorescence image). In each case, the correction was achieved after a maximum of 6 iterations with a gain of 0.6, which corresponds to a typical correction time of 500ms, mainly limited by the exposure time of the wavefront sensor.

**Figure 3:**
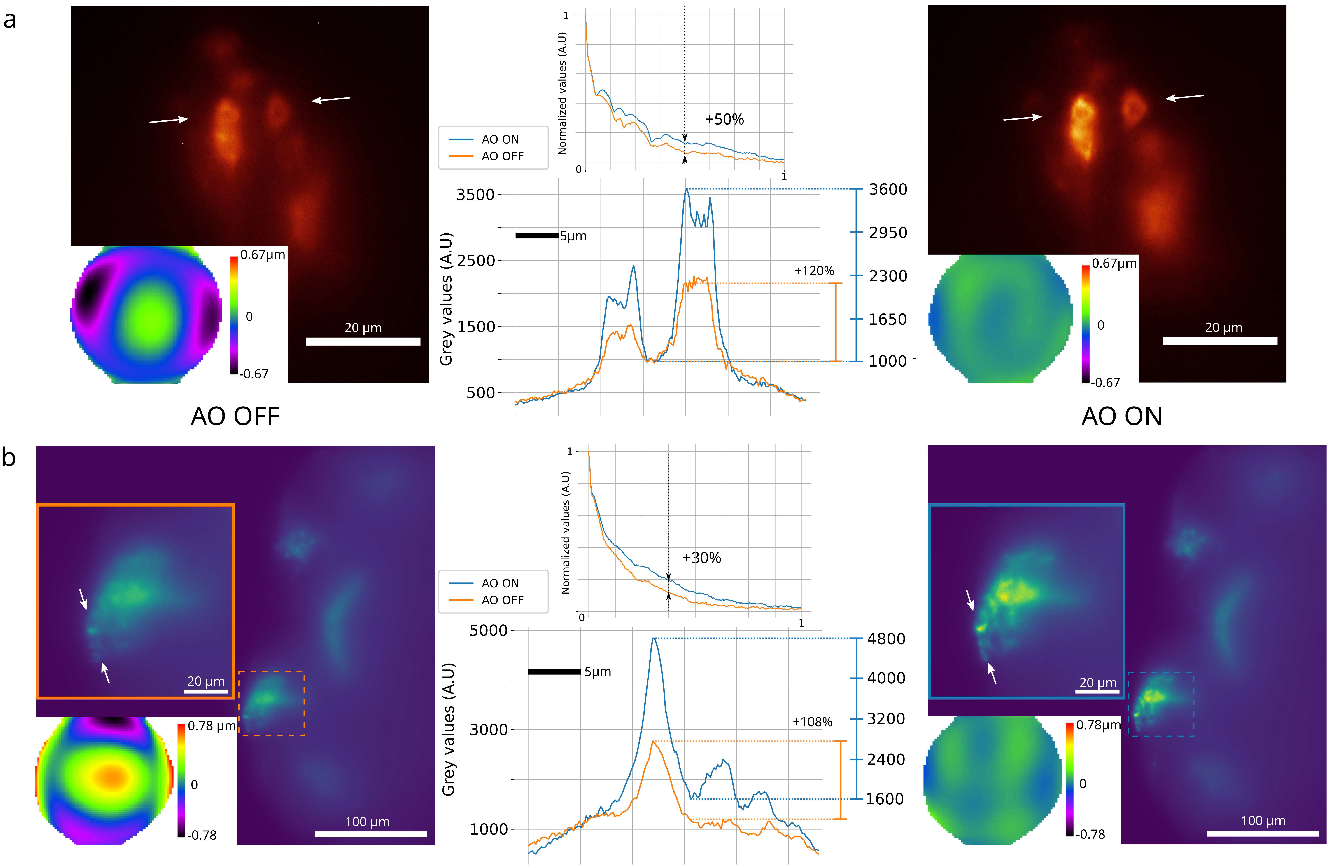
Image of freshly dissected *Drosophila* brains with FSB neurons expressing GFP and ChRFP, and associated measured wavefronts before and after AO correction expressed in μm. (a) cells bodies at an approximate depth of 30-55μm. (b) Full field of view of neuronal projections at an approximate depth of 60-80μm with a zoom on the corrected area. For (a) and (b), in the middle part, the top graph shows the average radial Fourier transform of the images before and after AO correction, and the bottom graph shows a profile corresponding to the line between white arrows before and after AO correction.

### 3.2 AO-LSFM for enhanced calcium neuroimaging in the live adult *Drosophila* brain

We then show the interest of our adaptive optics strategy using direct wavefront sensing using ESSH for calcium neuroimaging by demonstrating the gain in both sensitivity and spatial resolution in the images obtained with our AO-LSFM setup, at an approximate depth of 40μm in an *ex vivo Drosophila* brain. The fluorescence signal in the images in Fig.4 corresponds to emission of GCaMP7b that is proportionate to the native activity levels of sleep neurons at the time of the experiment [29, 30]. The fluorescence intensity levels recorded here are therefore consistent with those encountered in current functional neuroimaging experiments to map neuronal activity and ensure the relevance of this proof of concept. Exposure times of the imaging camera and the wavefront analyzer camera were respectively 100ms and 200ms. The wavefront is measured at 41μm (±10μm) in depth and the correction of the aberrations provides a gain in signal of a factor 2, clearly visible in the images of Fig.4 (a - top) and confirmed by the profiles on the two bright left neurons shown in Fig.4 (b - bottom). The signal enhancement can be observed at different depths (Fig.4 (a - bottom)), over a thickness of 10μm (see the volume reconstruction in Movie 1 in the supplementary material). This gain in intensity brought by AO directly corresponds to a gain of sensitivity of the imaging setup. Such a gain is of critical importance for functional experiments, since it provides the capability to detect and monitor neurons with lower activity, i.e. with faint signals close to the background level. An example of such neurons can be visualized in Fig.4 (b - top), as indicated by the red arrows. Before aberrations correction (Fig.4 (b top left), the intensity level of these neurons is of the same order of magnitude than the surrounding signal of brighter cells, whereas in Fig.4 (b top left), two neurons can be clearly identified thanks to the signal increase brought by AO. Zoom of the Fig.4 (b - middle) confirm the gain in sensitivity where AO allows to reveal low level signals of projections and highlight neuronal connections, indicated with red arrows and also observable in movie 1 (supplementary material). It has to be noted that the gain provided by AO is directly linked to the amount of aberrations arising from the sample. The experimental results from Fig.4 were obtained at an approximate depth of 40μm. As a consequence, it is expected that imaging deeper would provide a higher gain due to larger aberrations to be corrected, that might unfortunately be counterbalanced by scattering at some point, depending on the characteristics of the sample. In our experiments in the *Drosophila* brain, it was possible to demonstrate a significant gain from AO at depths down to 80μm for structural imaging and 40μm for calcium imaging. For now, we failed to report significant image improvement using AO at larger depths, which is attributed to scattering starting to be predominant.

**Figure 4:**
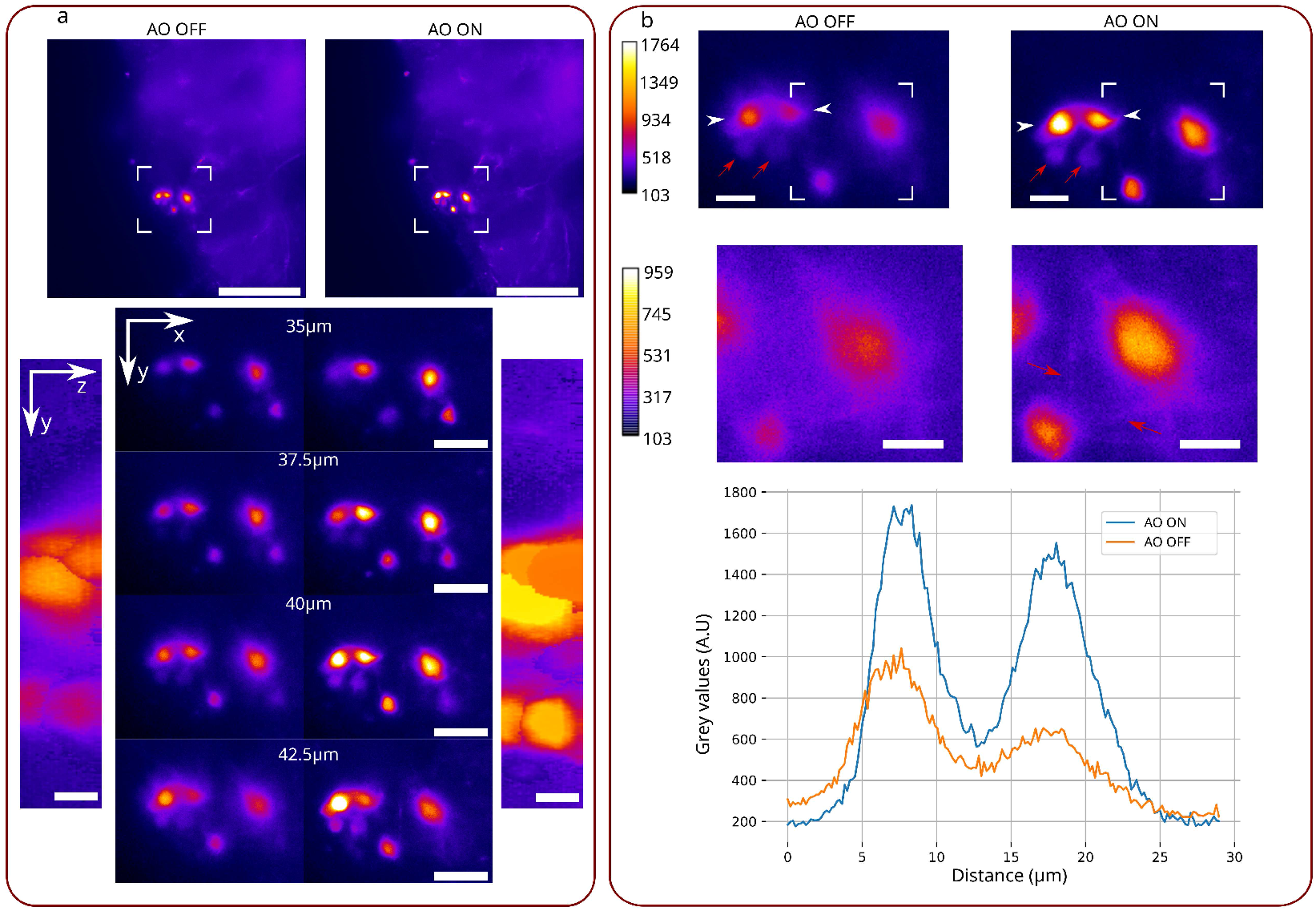
Neurons at 40μm (±10μm) deep labeled with GCaMP7b without and with adaptive optics. (a) (top) Full FOV image with delimited ROI. Scale bar: 100μm (bottom center) Zoom on previous ROI at several depths. Scale bar: 20μm. (bottom left and right) Max X projection of the stack without and with adaptive optics. Scale bar: 5μm (b) (top) Zoom on the ROI at z = 41μm (±10μm) with an adapted colorbar Scale bar: 10μm (center) Zoom on the delimited white rectangle above with a saturated colorbar (lower max value as compared to (top)), resulting in low level signals being more visible, such as neuronal connections indicated with red arrows (bottom) Profiles without (orange) and with (blue) adaptive optics taken between white arrows of top images.

## 4 Discussion

We developed an AO-LSFM system based on direct wavefront sensing not requiring any guide star and compatible with calcium neuroimaging. We demonstrated a typical 2-fold improvement in image contrast thanks to AO, with an optimal photon budget, which not requires the consumption of the photons used to build functional images. The approach requires a combination of a specific spectral excitation/detection and a dedicated sample preparation. Even though such sample preparation could be seen as a constraint, dual labelling enabling both structural and functional imaging is currently performed in routine in functional neuroimaging, where a stable fluorescence signal is needed as a reference. Indeed, the use of a second structural marker with a stable fluorescence intensity allows to normalize the activity levels of the different neurons in different samples in order to compare them over time without, for example, motion artifacts. This normalization is crucial during long-term recordings, and allows to easily localize and segment neurons as well as subcellular structures whose activity levels, and consequently the fluorescence signal, are of low intensity. Our approach enables fast AO correction at a current speed up to 10Hz, using a low complexity setup, demonstrating a significant speed increase, typically up to a factor of 5, as compared to our previous implementation [20], thanks to the reported dual labeling, photon-efficient strategy. One main advantage of AO based on direct wavefront sensing such as reported here is that it provides fast correction when compared to sensorless approaches, that rely on multiple image acquisitions, typically 20 to 40, for 3rd order Zernike modes [31], thus requiring 10s of seconds. This makes the device compatible with fast (500ms or less) AO correction, which is particularly beneficial to 3D imaging or imaging of dynamic processes requiring fast AO correction update. When targeting 3D imaging, depending on the extent of the axial isoplanetic patch, one or only a couple of AO loops can provide full correction over a large axial range, at the price of only one or several fast updates of the AO loop. One may object that reported imaging depths are limited. However, the adult *Drosophila* brain has been characterized as a strongly scattering sample when compared to other common animal models used in neuroimaging (e.g. Zebrafish embryo (*Danio rerio*), *C. Elegans*). Typically, in one-photon setups without AO, reported Signal-to-Backgroud ratio for this type of sample becomes weak starting at 40μm [32]. We report here structural and functional images down to 80μm and 40μm respectively with increased contrast. The ultimate challenge to image the whole adult *Drosophila* brain corresponds to an approximate depth of 200μm, with a good contrast over the whole volume allowing for the recognition of all cellular structures of interest and the detection of functional signals. Observing the activity of neurons in real time and how it is affected by other neurons and the environmental stimuli is essential to understand the functional organization and connection of the brain. The development of fluorescent sensors whose brightness varies according the activity level of the neurons has allowed scientists to use fluorescent microscopy to map large-scale neuronal activity in 3D with spatial resolution at the individual neuron level [4]. Nonetheless, the use of fluorescence intensity variation as a marker of cell activity has the inherent disadvantage of limited sensitivity which prevents the detection of low activity cells, leading to an unintentional detection bias towards active bright cells. Likewise, within the neuron itself, the cell body is usually easily detected, whereas projections with much weaker signals are much more difficult to observe. Moreover, the same neuron can transmit distinct information to several neurons via different branches of projections and synapses, with variable levels of activity, and thus variations in fluorescence intensity. Improving detection sensitivity is therefore crucial in order to be able to observe the neuronal activity at subcellular level and perform a local quantitative analysis [33]. Calcium imaging using GCaMP sensor, due to its fast kinetics, is used to study the fast-dynamic activity of the cell by tracing the changes in calcium traces over time in many publications (dF/F). For the specific case of the *Drosophila* line we studied, mainly involving neurons that induce sleep [34], the recording of dynamic activity over time requires *in vivo* imaging over a long time, including night time. As a first approach, taking into account current instrumental limits including software, and in order to minimize experimental constraints, we planned to demonstrate the gain brought by AO on calcium signal at a single time point. This approach was considered biologically relevant and informative since it is based on similar published studies in the *Drosophila* brain on sleep neurons facing the same experimental constraints. Indeed, similar to other fluorescent sensors such as CaLexA [35] and CaMPRI [36], in several publications using *Drosophila* flies, the fluorescent level of GCaMP for a certain time point is studied which is sometimes referred to as baseline fluorescent level, in other words an “activity snapshot” of the cell. This approach is used to compare the activity level between neurons in under different conditions such as feeding state [37] (Supp. Figure 7F and G) or at different time points of the day and after sleep deprivation [21] (Figure 5J) in order to link the baseline calcium level of the neurons and by consequences their activity levels to the experimental conditions. Thus, we believe the experiment using GCaMP7b sensor we performed in Fig. 4 reflects a functional imaging experiment that is on line with scientific publications targeting the same biology topic.

**Figure 5:**
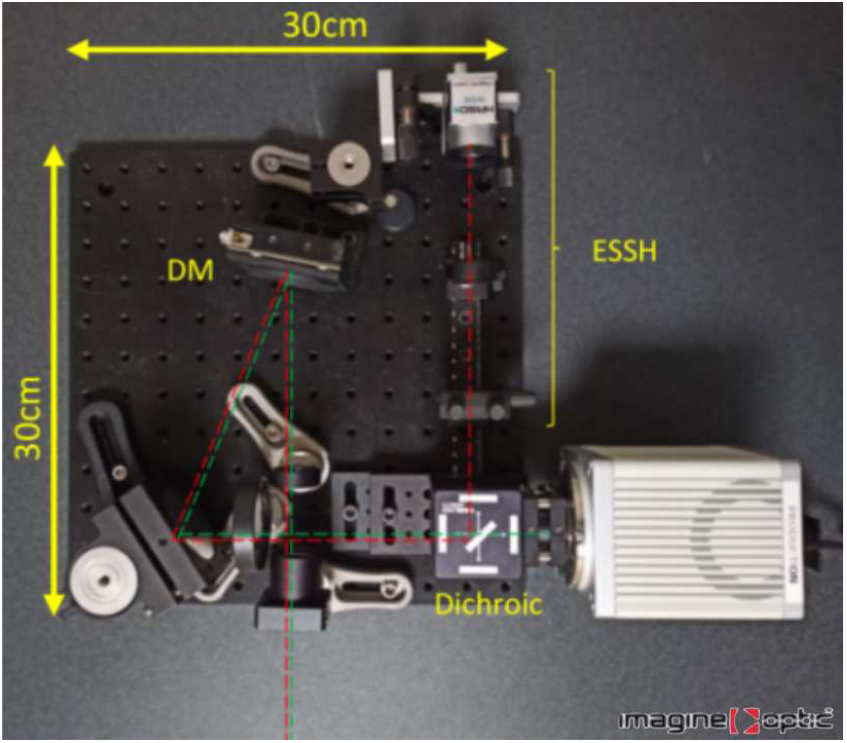
Compact prototype AO system for LSFM

The instrumental approach is compatible with the design of an add-on module for multiple geometries of LSFM setups, either research-oriented or commercial, since it only requires the insertion of the AO module before the imaging camera, such as for existing AO add-ons 1. A compact prototype module (30×30×10cm) has been designed as a first step toward a commercial AO add-on for LSFM systems, as illustrated in Fig 5. The prototype is currently being tested on a LSFM system based on an upright microscope frame and will be tested on several different LSFM configurations. On the current setup the light-sheet thickness is significantly larger than the depth of focus of the detection objective. The overall volume captured by each microlens of the ESSH wavefront sensor is thus 130×130×5μm^3^. As a consequence, the measured wavefront corresponds to an averaged wavefront from this volume, which might not be optimal if the size of the isoplanetic volumetric patch is significantly smaller. In our study, even if the 130×130μm^2^ FOV captured by each microlens of the ESSH is rather large, it has been shown that for our sample of interest the isoplanetic patch is at least of the same order of magnitude than this area. This is confirmed by the image obtained after AO correction of Fig. 3(b) where the wavefront is measured from the neurons and projections in the lower left area and an increase in fluorescence intensity can still be seen on the projections at the top of the image, as well as on Fig. S2 where a signal enhancement can be observed on the neuronal projections at the top of the image, at about 100μm far from the FOV of the analyzer. More generally, for samples whose isoplanatic patch is smaller than the analyzer FOV, the ESSH approach provides an interesting versatility, that has not been yet studied, for examples on more aberrated samples: it is possible to numerically adjust (reduce) the area over which intercorrelations are performed to measure wavefront slopes, which means that the size of the isoplanetic patch for a given sample can be characterized from a single acquisition, in order to define an optimal size for the computation of intercorrelation – as soon as such a size is smaller than the size of a microlens. Also, decreasing the light-sheet thickness would directly generate better scattering background rejection, suggesting significant room for even better image quality using our AO-LSFM. Finally, in our current system, AO is only implemented at the emission path: we are working on the introduction of optimization of the excitation light-sheet, such as a combination of “low-order AO” as reported in [38], with wavefront shaping in the excitation beam [39].

## Acknowledgements

We thank A. Fourgeaud (ESPCI) for providing custom mechanical parts and L. Bourdieu (ENS-IBENS) for fruitful discussion.

## Data availability

Data underlying the results presented in this paper are not publicly available at this time but may be obtained from the authors upon reasonable request.

## Author contributions

A.H, analysis, conceptualization, investigation, methodology, visualization, writing; G.F, investigation, methodology, sample’s preparation, writing; F.H conceptualization, funding acquisition, project administration, resources, supervision, writing; C.V, conceptualization, project administration; S.I, investigation; M.M, investigation; V.L, conceptualization, investigation; F.R, conceptualization, funding acquisition, project administration, resources, supervision; A.F, conceptualization, funding acquisition, project administration, resources, supervision, writing.

## Supplemental Materials

**Figure S1:**
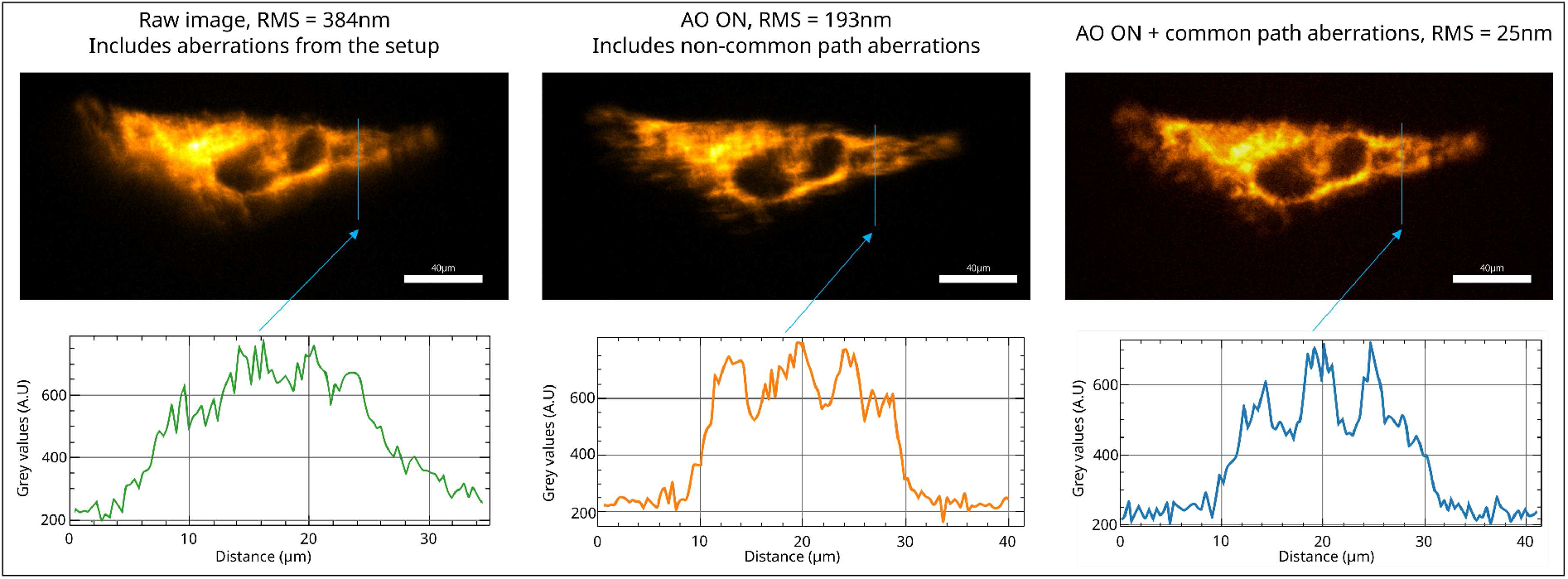
A living HeLa cell labeled with MitoTrackerGreen (Thermofisher) imaged using the same optical setup with an extra epifluorescence arm with a 488nm laser beam. The thickness of the cell is comparable to the thickness of the sheet in the light-sheet configuration and acts as a guide plane for the ESSH analyzer. Images on the scientific camera (left) with a static correction of the aberrations of the set-up from the objective to the wavefront analyzer (center) with a static correction of the set-up aberrations, including those of the non-common path to the imaging camera (right) with full correction of both set-up and sample induced aberrations.

**Figure S2:**
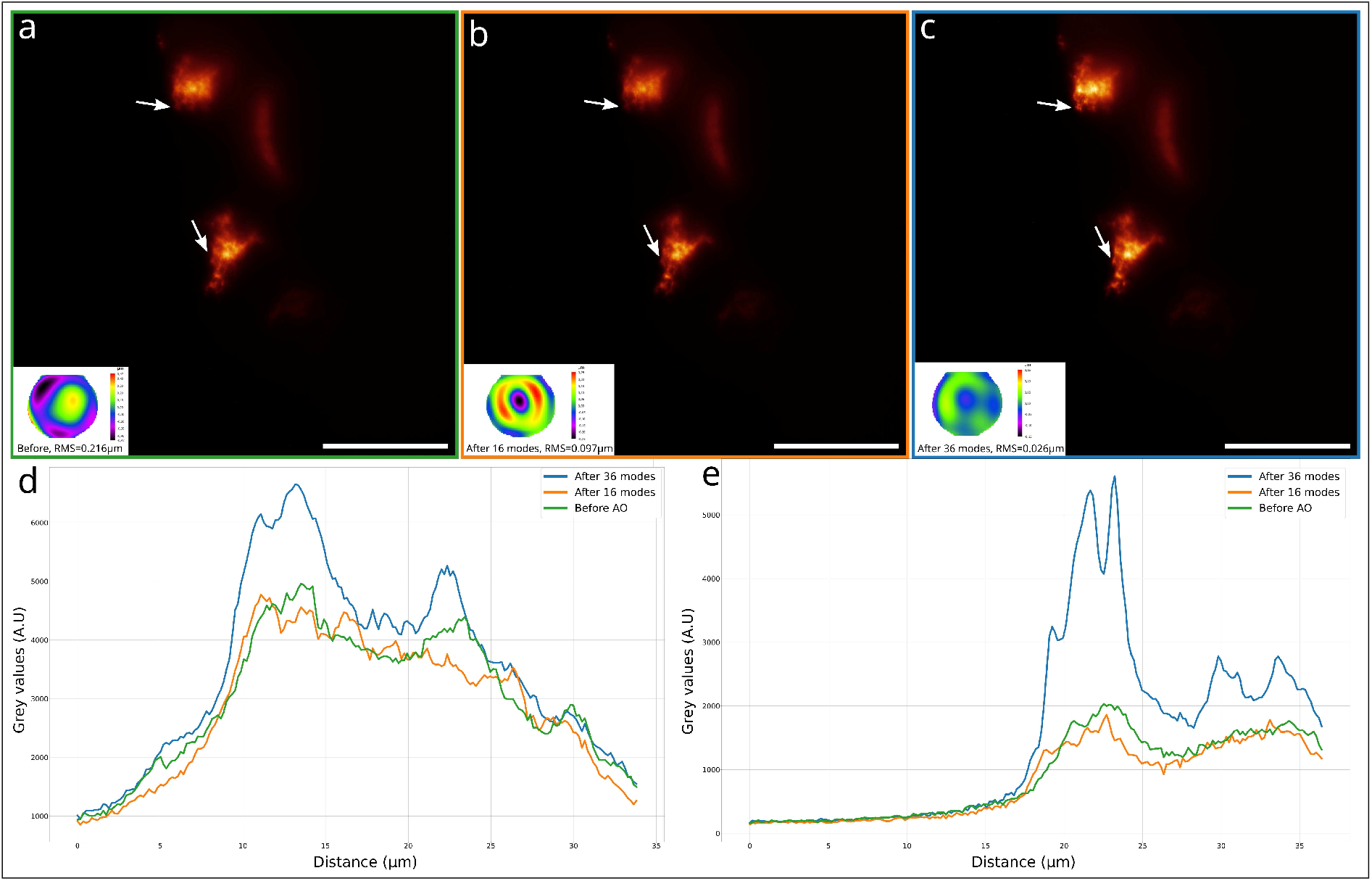
Full field of view of Fig.2 (top) projections of sleeping neurons at 60-80μm in depth, dual labeled with GFP/ChRFP, in an *ex vivo* adult *Drosophila* brain, with associated wavefronts and profiles in two separated areas (a) without correction, (b) with 16 DM modes for correction and (c) with 36 DM modes for correction. Line profiles in (d) correspond to centered white arrows and line profiles in (e) to top white arrows. The wavefront is estimated on a 130×130μm^2^ FOV located on the projections in the center of the image. Scale bar: 100μm

## Footnotes

1 https://www.imagine-optic.com/product/micao-3dsr/

